# Circadian and permanent pools of extracellular matrix co-exist in tendon tissue, but have distinct rates of turnover and differential responses to ageing

**DOI:** 10.1101/2024.08.09.607297

**Authors:** Anna Hoyle, Joan Chang, Marie FA Cutiongco, Ronan O’Cualain, Stacey Warwood, David Knight, Qing-Jun Meng, Karl E Kadler, Joe Swift

**Author notes:** Correspondence: QJM, KEK, and JS.

## Abstract

Heavy carbon isotopes in the tendons of people who grew up in the age of nuclear bomb testing have shown that the extracellular matrix (ECM), assembled during development, stays with us for life. However, recent work suggests that type-I collagen in ECM-rich mouse tendon exists in two pools: a permanent matrix, and a more soluble, circadian-regulated matrix. Despite this, the underlying regulation of such distinct pools is not understood. Here, we demonstrate using stable isotope labelling coupled with mass spectrometry proteomics that circadian and permanent matrix pools have significantly different half-lives. Furthermore, the properties of the matrix pools are altered during development and ageing. Tail tendon tissue was harvested from mice fed on a heavy-lysine diet; protein was then extracted for analysis using a sequential two-step protocol. The first, soluble fraction (‘F1’) was found to contain intracellular proteins, and a range of core and associated extracellular matrix proteins, including a pool of type-I collagen shown to be circadian-regulated. The remaining fraction (‘F2’) contained primarily collagens, including type-I collagen which did not show rhythmicity. In adult mice, matrix proteins extracted in the F1 pool had significantly shorter half-lives than F2, including type-I collagen which had half-lives of 4 ± 2 days in F1, compared to 700 ± 100 days in F2. Circadian-regulated matrix proteins were found to have significantly faster turnover than non-circadian in adult mice, but this distinction was lost in older animals. This work identifies protein turnover as the underlying mechanism for the circadian/permanent model of tendon matrix, and suggests a loss of circadian regulation as a characteristic of ECM ageing.

## INTRODUCTION

Protein turnover is a crucial mechanism in the maintenance of tissue homeostasis. While the majority of intracellular proteins are replaced on timescales of hours to days (1), the extracellular matrix (ECM) is subject to slower remodelling and is therefore more susceptible to accumulation of damage (2). The ECM consists of a complex network of proteins including core, structural proteins such as collagens, proteoglycans, and glycoproteins, as well as many ECM-associated proteins which bind to and regulate it (3). Collagen is the most prevalent protein in humans, with the bulk of it thought to be assembled during development to form permanent structures that last throughout our lifetimes (4, 5). This ‘minimal turnover’ model has been evidenced by tracking the heavy carbon isotopes incorporated into tissues following atmospheric nuclear bomb testing, and analysis of advanced glycation endpoint (AGE) modifications, racemization and labelled amino acids in ECM proteins (5–8). However, the minimal turnover model is challenged by evidence that type-I collagen is produced in response to exercise and mechanical loading (9, 10). These seemingly contradictory observations can be rationalised by a model in which ECM components coexist in multiple states within the same tissue (5); this has been supported by recent work showing that type-I collagens (Col1a1 and Col1a2) in mouse tendon exist in two pools: a permanent matrix, and a more soluble matrix that is regulated over a circadian cycle (11).

Circadian rhythms are essential for cellular and tissue homeostasis – they enable organisms to anticipate daily changes in their environment and respond accordingly. Although synchronised by the suprachiasmatic nucleus in the hypothalamus, almost all tissues, including the tendon, contain a peripheral circadian clock which regulates localised circadian functions (11). Peripheral clocks have been found to filter circadian signals from the brain to optimise tissue-specific functioning, including ECM homeostasis, in tissue such as muscle and skin (12). Support for a permanent/circadian matrix model can be found in studies showing circadian rhythmicity in levels of type-I collagen: in humans, precursors of mature type-I collagen (propeptides) were found to be rhythmic in serum (13), while collagen degradation products were rhythmic in urine (14). Rhythmic collagen expression previously characterised in mouse tendon tissue (11) has now also been demonstrated in human tendons (15). Patients with chronic tendinopathy, long-term tendon inflammation and damage had dampened circadian rhythms with decreased type-I collagen production overnight. Other work has shown that mice with disrupted circadian rhythms had abnormalities in collagen-rich tissues (16, 17). These findings demonstrate the importance of understanding the complexities of ECM turnover and suggest the circadian clock as a key modulator of ECM homeostasis and a potential therapeutic target. Whilst circadian rhythmicity of ECM components has been characterised in a range of tissues, including kidney (18), cartilage (19) and intervertebral disc (20), our understanding of the underlying regulatory mechanisms remains limited. Insight can be gained from the findings of Chang *et al.*, specifically that analysis of a more soluble fraction of the ECM reveals a complexity of regulation that is masked in bulk analysis (11). Variation in protein turnover rate as a function of solubility or localisation has previously been shown in tendon (21), cartilage (22, 23), liver (24) and lung (25).

Ageing is associated with a loss of ECM integrity and structural changes resulting from altered dynamics and aggregation of damage. Changes to the ECM and its regulation are linked to a number of the systematic ‘hallmarks of ageing’ (26), as well as tissue-specific alterations. An understanding of age-associated changes in the ECM at a tissue-specific level will be vital to addressing pathologies in an ageing population, particularly those related to impairment of mechanical function. Tendon tissue stiffens with age, accompanied by an accumulation of damage, leading to reduced mechanical resilience and an increased risk of injury (27, 28). Aging is also associated with a suppressed cellular response to the mechanical environment (29, 30) and dysregulation of the circadian clock (31). Defects in proteinases targeting type-I collagen and collagenase-resistant mutations to collagen both lead to signs of premature aging, indicating the importance of maintained ECM degradation pathways (32, 33). Additionally, aged tendons display altered distributions of dimensions in their collagen fibrils, an observation also made in the tendons of mice with knockout of decorin, a proteoglycan important in ECM assembly (34). Studies of remodelling in heart tissue have shown that measures of protein turnover can provide more insight than protein abundance alone (35). Furthermore, recent work has shown that coordinated circadian regulation of protein synthesis and degradation can result in rhythmic turnover, rather than cycling of abundance (36).

Despite evidence of the importance of maintained ECM remodelling (i.e., balancing processes of matrix deposition and degradation), studies of protein turnover have remained limited. Seminal work tracking the rate of incorporation of ^14^C into tissues following atmospheric testing of nuclear weapons between 1955 and 1963 established very low rates of carbon turnover in bulk human tendon (5), eye lens (37), and dental enamel (38), relative to tissues such as skeletal muscle. Other methods for estimating the longevity of proteins within tissues have tracked post translational modifications (PTMs), although these can be confounded by factors such as temperature and states of disease (39, 40). Developments in mass spectrometry (MS) proteomics have enabled estimates of protein turnover rate by examining how stable heavy isotopes are incorporated; importantly, these MS methods can identify specifically which proteins have incorporated the heavy labels. Labelling with deuterated water was used to study protein turnover in rat Achilles tendon, establishing rates specific to different regions of the tissue (21). Introduction of heavy amino acids, specifically ^13^C-lysine, into animal feed has allowed development of heavy-labelled mouse models. These were initially used as standards for quantitative MS, but have since been adopted as tools to study protein turnover (41–43). Recent work has used this approach to study ECM turnover in bone and cartilage, demonstrating a clear decline in matrix synthesis with age (44).

In this study, we used MS to examine tendon tissues from mice fed a ^13^C-supplemented diet. Subsequent bioinformatic analysis of the rates of incorporation of the heavy-label allowed evaluation of the half-lives of 1790 proteins. By extracting and analysing proteins from the circadian matrix pool, as in Chang *et al.* (11), in addition to analysis of the sparingly soluble non-circadian pool, we demonstrated significantly different rates of protein turnover. By repeating measurements in young and old mice, we found significantly altered ECM dynamics in ageing. This analysis has also established decreasing distinction between the turnover characteristics of circadian/soluble and non-circadian/permanent matrix pools, suggestive of a loss of circadian regulation in ageing.

## RESULTS

### Two-stage solubilisation of protein from mouse tail tendon reveals circadian and non-circadian pools of extracellular matrix

Transmission electron microscopy (TEM) images of cross-sections through adult mouse tail tendon tissue, orientated perpendicular to the direction of mechanical loading, have previously identified type-I collagen fibrils with a distribution of morphologies (Fig. 1A) (11). To aid characterisation of this distribution, fibrils were categorised as ‘D1’, with diameters < 75 nm; ‘D2’, diameters 75 – 150 nm; and ‘D3’, diameters > 150 nm (45). The distributions of fibre morphologies have been shown to vary over a circadian (24-hour) cycle, with TEM images of tendons analysed at 12-hour intervals (from populations of circadian-synchronised mice) exhibiting cyclic loss and recovery of D1 fibrils (Fig. 1B) (11). A mild detergent wash of mouse tendon tissue (fraction 1, ‘F1’) acquired from a population of mice entrained with 12 hr/12 hr light/dark cycles was demonstrated to extract proteins with significant circadian rhythmicity over a period of 48 hours (11). Rhythmic proteins included the type-I collagens Col1a1 and Col1a2 (quantified by MS proteomics; *q*-values 0.007 and 0.014, respectively). We have now shown that a subsequent high salt extraction of the same material (fraction 2, ‘F2’) resulted in solubilisation of the pellet, but that proteomic analysis of F2 no longer showed circadian rhythmicity in type-I collagen levels (*q*-values = 1; Fig. 1C). Analysis of tendon cross-sections by TEM showed that the F1 extraction preferentially solubilised D1 fibrils, meaning that F2 contained primarily D2 and D3 material (Supplementary Fig. S1).

**Figure 1.**
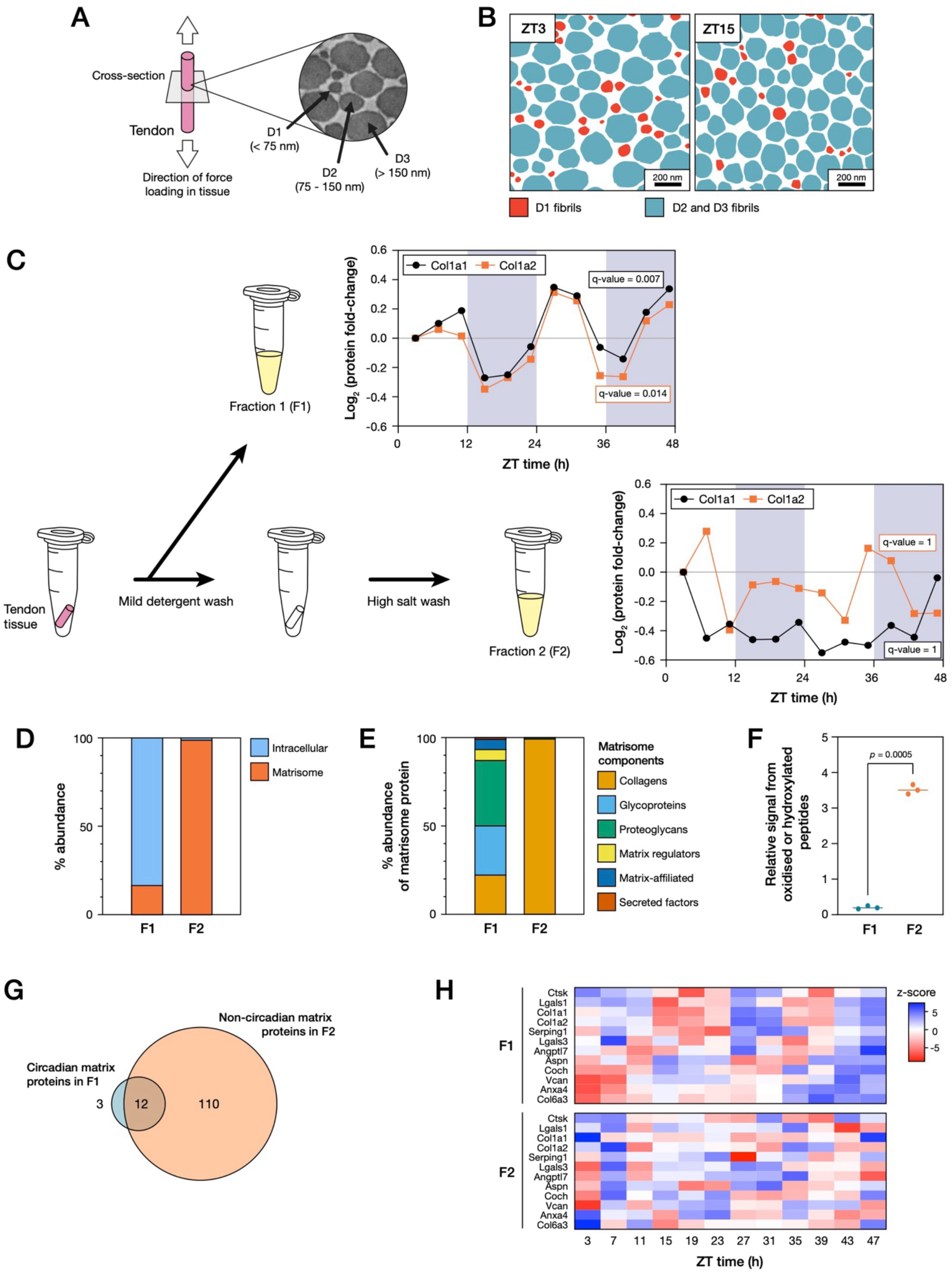
Identification of circadian and non-circadian matrix pools in mouse tendon tissue. (**A**) Schematic showing how mouse tail tendon sections prepared for transmission electron microscopy (TEM) were taken perpendicular to the length of the tendon, its constitutive fibrils, and the direction of mechanical loading. Fibrils were classified by their diameters: ‘D1’, < 75 nm; ‘D2’, 75 – 150 nm; ‘D3’, > 150 nm. (**B**) Cartoon illustrating analysis of tail tendon fibril morphologies in rhythm-synchronised mice at a 12-hour interval (zeitgeber time, ZT; ZT3, 3 hours after light-on; ZT15, 3 hours after light-off), showing loss of D1 fibrils. Figure panels (A) and (B) adapted with permission from Chang *at al.* (11). (**C**) Schematic showing the two-step solubilisation protocol used for protein extraction from mouse tendon tissue. Plots show changes in relative abundances of Col1a1 and Col1a2 over 48 hours, measured by mass spectrometry (MS) following extraction in a mild detergent wash of the tendon tissue (fraction 1, ‘F1’) and a subsequent harsh solubilisation protocol applied to the same material (fraction 2, ‘F2’). (**D**) Fraction of the total signal detected by MS from peptides derived of intracellular vs. matrisome proteins in F1 and F2. (**E**) Distribution of the signal derived from matrisome proteins between the structural and functional classifications of the matrisome in F1 and F2. Matrisome proteins and classifications in (D) and (E) are described by Shao *et al.* (46). (**F**) Relative signal from peptides detected by MS with oxidation or hydroxylation modifications in F1 and F2; *p*-value determined by t-test. (**G**) Of 125 matrisome proteins identified in mouse tail tendon tissue, 122 were found to lack significant circadian rhythmicity in F2 (*q*-value > 0.05). However, of these, 12 proteins had shown rhythmicity in F1. (**H**) Heatmap showing matrix proteins with significant circadian rhythmicity in F1, but that lack rhythmicity in F2. Blue and red depict up and down regulation, respectively. Proteomics analysis from *n* = 4 animals per time point.

Proteins identified in MS analysis of F1 and F2 fractions were compared to the matrisome database of recognised ECM components (MatrisomeDB, 46). F1 was found to contain primarily intracellular material, along with ∼20% extracellular material (determined from the fraction of total peptide signal in MS, as a surrogate measure for relative concentration); F2 by contrast contained almost exclusively extracellular proteins (Fig. 1D). Analysis of the matrisome has been used to classify ECM components by structural and functional features (46). In the F1 fraction, we found an abundance of signal from protein-derived peptides covering a number of matrisome classifications: primarily glycoproteins and proteoglycans, but also collagens, matrix regulators and matrix-affiliated proteins; the majority of signal in the F2 fraction came from collagen-derived peptides (Fig. 1E). Consistent with a greater proportion of extracellular material, the fraction of peptides detected in their oxidised states was significantly greater in F2 than in F1 (*p*-value = 0.0005; Fig. 1F). Levels of a combined set of 125 ECM proteins detected in both F1 and F2 fractions were analysed over 48 hours to determine significant circadian rhythmicity (analysis at 4-hour intervals; *n* = 4 animals per timepoint; *q*-values < 0.05). Of 15 proteins found to have significant circadian rhythmicity in F1 (11), 12 did not exhibit rhythmicity in F2 (Fig. 1G). This suggests a set of ECM proteins – including primary connective tissue constituents Col1a1 and Col1a2 – that simultaneously exist in both circadian/soluble and non-circadian/permanent states within the same tissue (Fig. 1H).

### Heavy isotope labelling allows estimation of rates of turnover of intra- and extracellular proteins in mouse tail tendon

By measuring the rate at which heavy-labelled amino acids are incorporated into newly synthesised proteins (Fig. 2A), methods such as stable isotope labelling with amino acids in cell culture (SILAC) (47) can be used to establish protein translation rates (48), and have been adapted to work in mouse models (41, 44). Peptides derived from biochemically identical ‘heavy’ and ‘light’ versions of the same protein can be distinguished by MS, allowing pre-existing and newly synthesised (post-labelling) proteins to be independently quantified. C57BL/6 mice were fed a heavy-labelled amino acid (^13^C-lysine) diet for either 1, 2, 4 or 8 weeks alongside control mice fed on an equivalent ^12^C-lysine diet (Fig. 2B). Mice were sacrificed at 22 weeks and tail tendon tissue was excised, solubilized in F1 (circadian/soluble) and F2 (non-circadian/permanent) fractions, and analysed by MS (*n* = 3 animals per timepoint). The quantity of ^13^C-lysine incorporated into proteins was found to increase significantly as animals were fed the heavy diet for longer periods of time (*p* < 0.0001; Supplementary Fig. S2A), and principal component analysis (PCA) showed clustering of peptide-level MS data by heavy diet feeding duration in both F1 and F2 fractions (Supplementary Figs. S2B, C). The heavy label was found to be incorporated into proteins in the F1 fraction at a significantly greater rate (Supplementary Fig. S2D), consistent with this fraction containing a greater proportion of shorter-lived intracellular proteins, relative to the primarily extracellular constitution of F2 (Fig. 1D).

**Figure 2.**
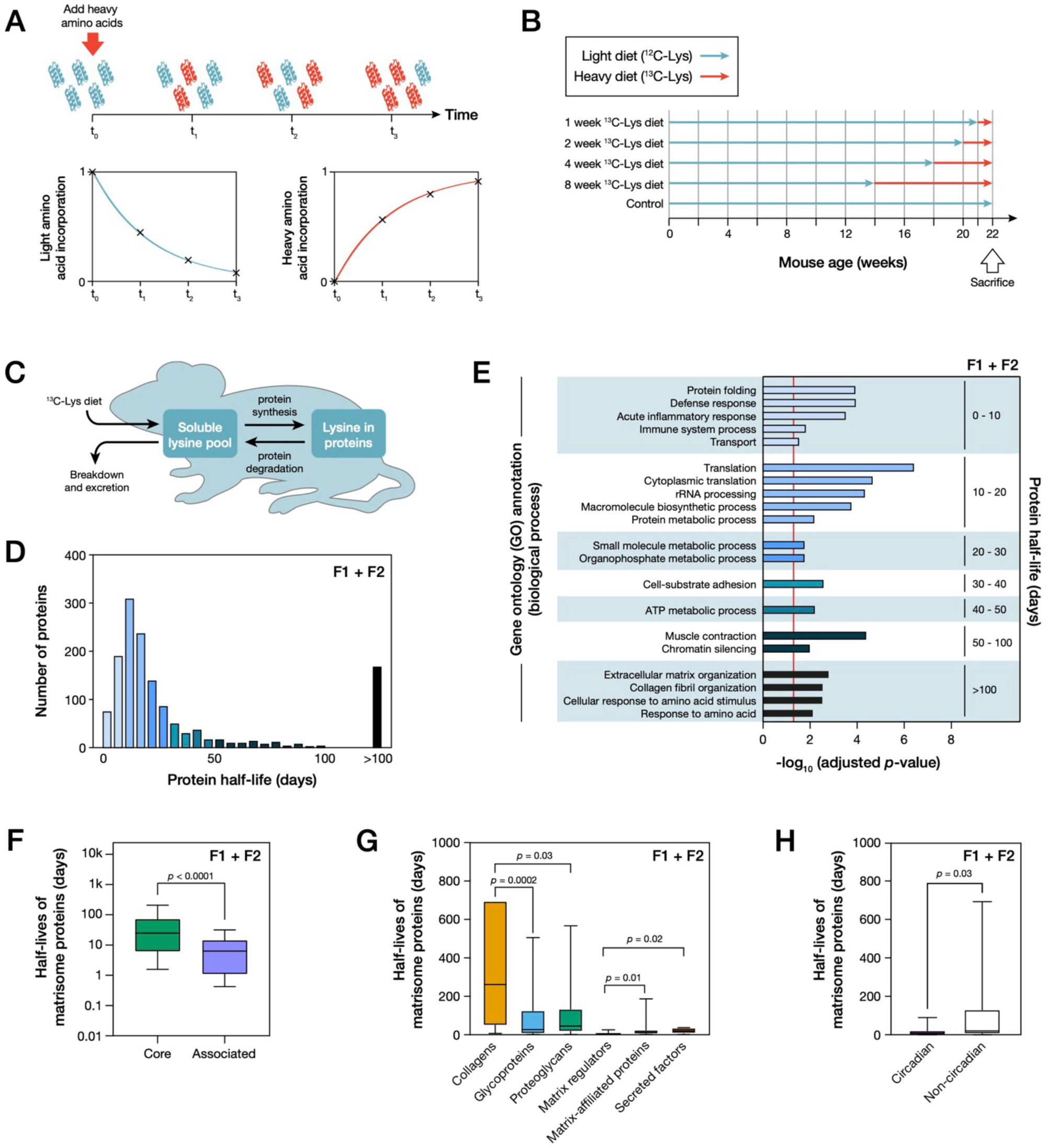
Heavy isotope labelling identifies relationships between protein function and half-life in mouse tail tendon tissue. (**A**) Schematic showing the principle of heavy isotope labelling: following addition of heavy-isotope variants of amino acids, metabolism in cells or organisms means that, over time, ‘light’ versions of proteins (blue) are replaced by chemically indistinguishable ‘heavy’ versions (orange). Peptides derived from heavy and light proteins can be independently quantified by mass spectrometry (MS). (**B**) Schematic showing the ‘light diet’ (containing ^12^C-lysine amino acids) vs. ‘heavy diet’ (^13^C-lysine) mouse feeding regimes used in this study. Following sacrifice, tail tendon tissue was fractionated into circadian/soluble (F1) and non-circadian/permanent (F2) fractions for analysis by MS. (**C**) Schematic showing the fate of ^13^C-lysine in mice, forming the basis of models for protein half-life estimation described by Alevra *et al.* (49). (**D**) Combined distribution of calculated protein half-lives in both F1 and F2 fractions of mouse tail tendon tissue. (**E**) Biological process gene ontology (GO) term enrichment within ranges of protein half-lives shown in (D). Red line indicates an adjusted *p*-value = 0.05. (**F**) Distribution of half-lives of ECM proteins in adult mouse tail tendon tissue in ‘core’ and ‘matrix-associated’ matrisome classifications; *p*-value from Mann-Whitney test. (**G**) Distribution of half-lives of ECM proteins in adult mouse tail tendon tissue within matrisome subclasses; *p*-values determined by Kruskal-Wallis test. (**H**) Non-circadian matrisome proteins had significantly longer half-lives than those with circadian rhythmicity; *p*-value from Mann-Whitney test. Matrisome structure/function classifications were described by Shao *et al.* (46); circadian matrisome proteins were identified by Chang *et al.* (11). Figure panels (D)-(H) combine data from both F1 and F2 fractions; box-whisker plots indicate medians, 10^th^ and 90^th^ percentiles, and minimum/maximum values. All data derived from proteomics analysis with *n* = 3 animals per condition; protein half-lives determined using models described by Alevra *et al.* (49).

By accounting for the presence of a ‘reservoir’ of free lysine within our labelled animals (Fig. 2C), we converted the ratios between MS signal from heavy and light peptides (‘H/L ratios’) into estimates of protein half-life, as described by Alevra *et al.* (49). Half-lives were significantly greater in F2 than in F1 (*p* < 0.0001; Supplementary Fig. S2E). A histogram of all protein turnover rates calculated from a combined analysis of F1 and F2 fractions showed a peak in the distribution of half-lives at around 10 days (Fig. 2D), consistent with previous reports of half-lives in the order of days for intracellular proteins in fibroblasts cultured *in vitro* (1). However, we also discovered a substantial number of longer-lived proteins, with half-lives exceeding 100 days. Analysis of enrichment of biological process gene ontology (GO) annotations within protein groups determined by half-life (Fig. 2E) showed the shortest-lived proteins to be associated with immune response, and protein metabolism, folding and transport. Proteins with intermediate and longer half-lives were related to functions of adenosine triphosphate (ATP) metabolism, cell-substrate adhesion (i.e., regulating cellular interactions with the ECM), and chromatin silencing. The longest-lived proteins were associated with ECM and collagen fibril organisation.

To look more specifically at matrix proteins within our data, protein half-lives in fractions F1 and F2 were classified according to annotations in MatrisomeDB (46). Core matrisome proteins were found to have significantly longer half-lives than matrisome-associated components (*p* < 0.0001; Fig. 2F), perhaps indicative of a difference between primarily structural versus signalling and regulatory roles. Within the core matrisome group, collagens were found to have significantly longer half-lives than either glycoproteins or proteoglycans; matrix regulators were also significantly longer lived than matrix-affiliated proteins and secreted factors (Fig. 2G). Importantly, we found that matrisome proteins that have previously been shown to exhibit circadian rhythmicity (11) had significantly shorter half-lives than non-circadian matrisome proteins (*p* = 0.03; Fig. 2H), suggesting a requirement for dynamic regulation for at least a subpopulation of these proteins.

### Circadian/soluble and non-circadian/permanent pools of extracellular matrix proteins in mouse tail tendon have distinct rates of turnover

We hypothesised that a principal biochemical difference between two co-existing pools of the same ECM constituents would be their respective rates of turnover. In order to test this, we compared the rates of incorporation of ^13^C-lysine into matrix proteins in F1 and F2 fractions. We found that H/L ratios from matrisome-derived peptides increased as mice were fed the heavy label diet for longer periods, but that the extent of label incorporation was significantly lower in F2 versus F1 (Fig. 3A). This was reflected in protein half-lives, with collagen, glycoprotein and matrix regulator matrisome groups all significantly longer lived in F2 than F1 (Fig. 3B). By selecting matrisome proteins from F2 with half-lives greater than a year, we used the STRING database of known protein-protein interactions (50) to produce a network diagram of interacting proteins within a ‘permanent matrix’ (Fig. 3C). The most abundant proteins in the permanent matrix, as determined by MS signal intensity, were Col1a1 and Col1a2, but type II, IV, V, VI, VII and XI collagens, glycoproteins such as fibrillin-1 (Fbn1) and proteoglycans were also included. 15 matrisome proteins were found to be turned over at a significantly greater rate in F1 versus F2 (*p*-values < 0.05; Fig. 3D), 11 of which were associated with the core matrisome. Col1a1, Col1a2 and the proteoglycan asporin (Aspn) were found to have significant differences in half-life between F1 and F2 pools, as well as to exhibit circadian rhythmicity in F1, but not in F2 (Fig. 1H).

**Figure 3.**
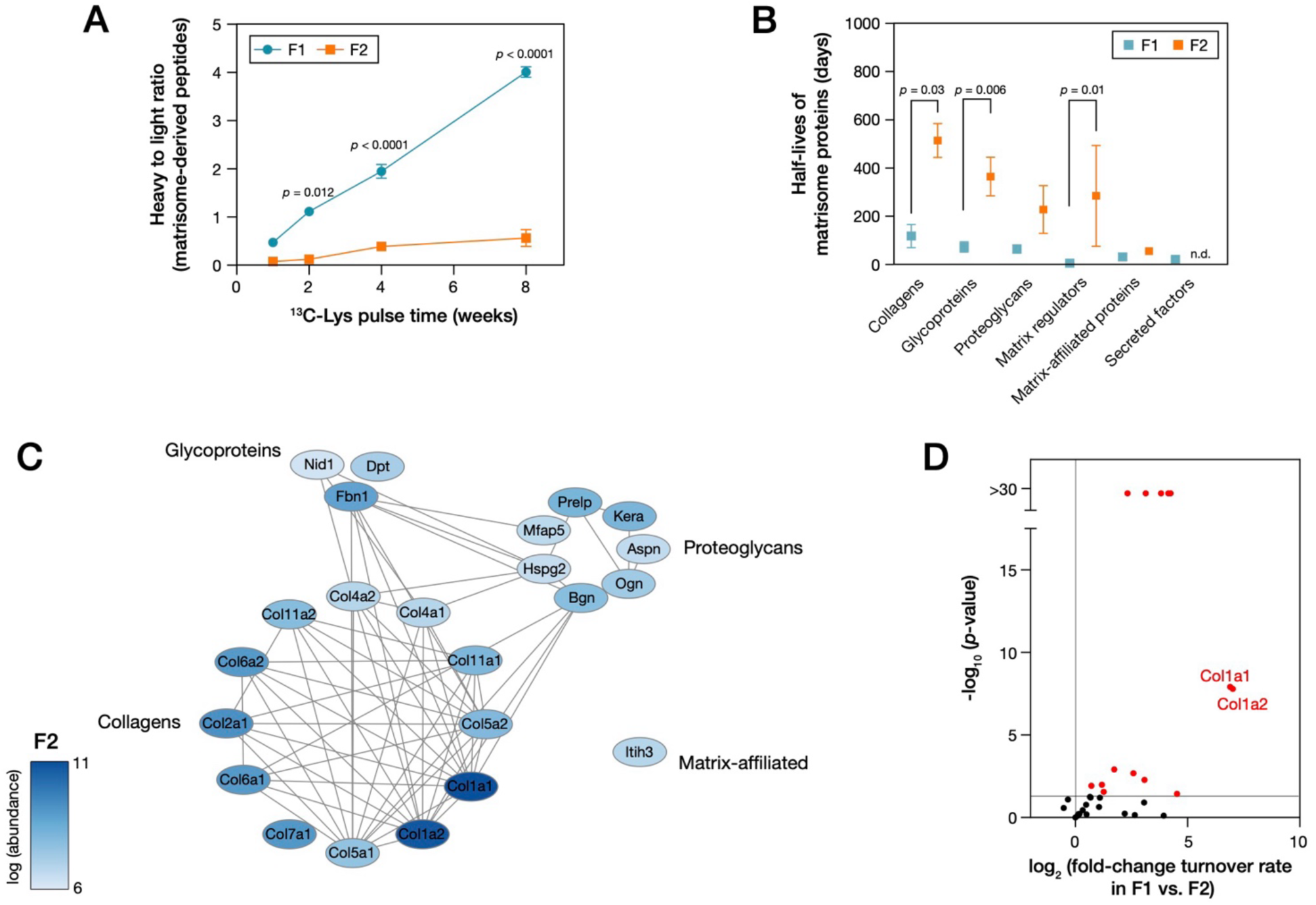
Mouse tail tendon tissue contains two pools of extracellular matrix (ECM) protein with significantly different rates of turnover. (**A**) Ratios of signal from heavy and light peptides derived from the matrisome in analysis of F1 and F2 fractions of tail tendon from 22 week old mice fed a ^13^C-Lys diet for 1, 2, 4 or 8 weeks. Points indicate mean ± s.e.m.; *p*-values from two-way ANOVA. (**B**) Average half-lives of proteins in matrisome structure/function classifications, comparing F1 to F2 fractions. No half-lives were determined for secreted factor proteins in F2 (n.d. = ‘not detected’); *p*-values from Kruskal-Wallis tests. (**C**) Network of interactions between ECM proteins detected in F2 with a half-life measured to be greater than a year. Connections are interactions identified in the STRING database with an interaction score > 0.7 (50). Nodes are proteins coloured by their relative abundance identified in tendon. (**D**) Volcano plot showing the difference in matrisome protein turnover rates in F1 vs. F2 fractions. All data derived from proteomics analysis with *n* = 3 animals per condition; protein half-lives determined using models described by Alevra *et al.* (49).

Examining type-I collagens – the major protein constituents of tendon tissue – in greater detail, we found that the H/L ratios from collagen-derived peptides were significantly greater in F1 versus F2 pools at all labelling timepoints (*p*-values < 0.0001; Fig. 4A). This data allowed us to calculate half-lives of Col1a1 and Col1a2 as 4 ± 2 days in F1, and 700 ± 100 days in F2. In order to investigate whether this separation in half-lives reflected mature collagens, or potentially a pool of intracellular procollagen precursors, we examined if these proteins had been post-translationally processed. Type-I collagens are subject to a range of PTMs, including hydroxylation of proline and lysine, and glycosylation of hydroxylated lysine; each of these occur enzymatically within the cell prior to secretion (51). N- and C-termini of procollagen triple helices are enzymatically cleaved off, resulting in mature collagen, which can subsequently assemble into higher-order fibrillar structures. H/L ratios from Col1a1- and Col1a2-derived peptides were mapped onto the protein sequences, with peptides found to originate from the main chain region, the C-propeptide region, and the site of C-proteinase cleavage (Fig. 4B). Peptides corresponding to the propeptide region and cleavage site showed significantly faster turnover than the main chain region in both F1 and F2 fractions, and in Col1a1 and Col1a2 (Figs. 4C, D). If a sample contained only the procollagen, all regions of the protein would be turned over at the same rate. We therefore conclude that both F1 and F2 fractions contained precursor and mature forms of type-I collagen.

**Figure 4.**
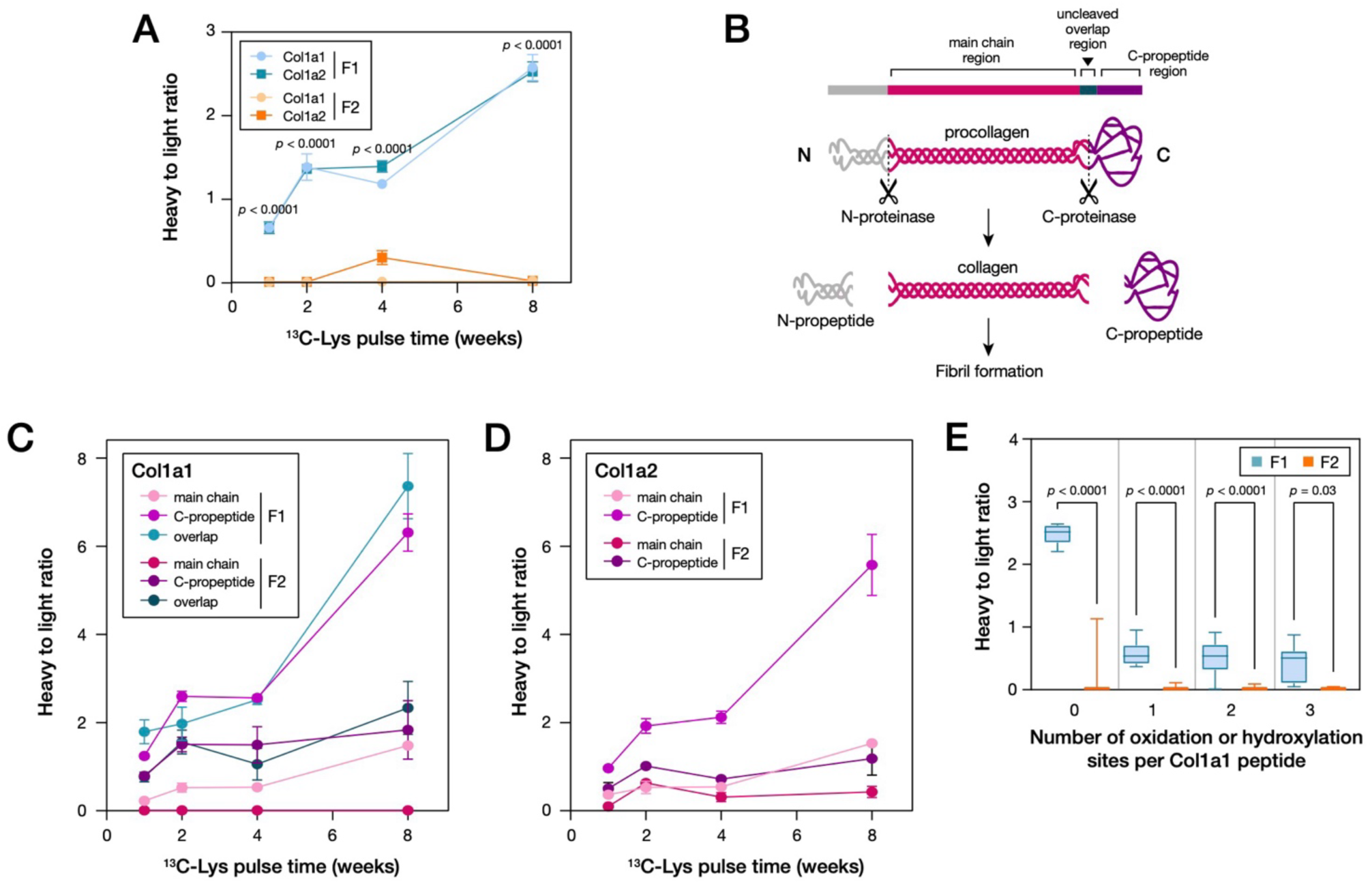
Type-I collagen has a significantly longer half-life in F2 vs. F1, but both pools are subject to post-translational processing. (**A**) Average ratios of signal from heavy and light peptides derived from Col1a1 and Col1a2 in analysis of F1 and F2 fractions of tail tendon from 22 week old mice fed a ^13^C-Lys diet for 1, 2, 4 or 8 weeks. Points indicate mean ± s.e.m.; *p*-values from two-way ANOVA. (**B**) Schematic showing type-I collagen processing. Adapted with permission from Canty and Kadler (52). (**C**) Average heavy to light ratios of signal from peptides originating from different regions of the Col1a1 sequence in fractions F1 and F2. (**D**) Average heavy to light ratios of signal from peptides originating from different regions of the Col1a2 sequence in fractions F1 and F2 (note that peptides from the overlap region were not detected in Col1a2). (**E**) Average ratios of signal from heavy and light peptides derived from Col1a1, in 22 week old mice fed a ^13^C-Lys diet for 4 weeks, as a function of the number of oxidation or hydroxylation modifications per peptide. *P*-values from Kruskal-Wallis tests. All data derived from proteomics analysis with *n* = 3 animals per condition.

Collagens may be subjected to further oxidation and crosslinking in the extracellular space, which can occur spontaneously or through enzymatic action (51, 52). Our MS analysis allowed quantification of oxidation and hydroxylation by searching for a mass increase of +15.955 Da per modification, although it could not distinguish between the two. By considering H/L ratios of modified and unmodified peptides, we determined that Col1a1 was turned over significantly faster in F1 versus F2 irrespective of the number of oxidation/hydroxylation modifications (Fig. 4E), but that oxidised/hydroxylated protein was significantly more stable than the unmodified forms in both F1 and F2 (Supplementary Figs. S3A, B). This result suggests correlation between the persistence of type-I collagen and the extent of its oxidation/hydroxylation. Oxidation can be a precursor to the formation of covalent crosslinks between collagen molecules. Although these modifications are difficult to observe directly by MS, their effects may be seen by a reduction in protein solubility. Here we note that although in MS sample preparation our homogenisation and solubilization protocol was optimised to the extent that no visible insoluble pellet was observed, some fraction of heavily crosslinked ECM protein may have remained undetectable; the consequence of this would be an underestimation of the half-lives of type-I collagens in the non-circadian/permanent pool.

### The composition of mouse tail tendon tissue changes in development and ageing

To investigate the effects of ageing on tail tendon tissue, groups of mice were fed a ^13^C-lysine diet for 4 weeks prior to sacrifice at either 8, 22, 52 or 78 weeks of age; these groups will hereafter be referred to as ‘young’, ‘adult’, ‘old’ and ‘aged’, respectively (Fig. 5A). As previously, tissue was analysed by MS in F1 and F2 fractions. Before calculating protein half-lives, we considered changes to the bulk composition of the tissue. Analysis of enrichment of biological process GO annotations in old and aged versus adult mice showed significant down regulation of processes associated with protein metabolism (translation and the ribosome), ECM regulation (collagen fibril organisation) and cellular mechanosensing (actin nucleation, integrin signalling and cell adhesion) (*p* < 0.05, Supplementary Fig. S4A). These changes were consistent with those observed in earlier proteomic studies of cellular senescence (29, 53). Within the more soluble F1 fraction, the contribution of matrisome components to the total peptide signal increased between young and adult, before decreasing in the old and aged mice (Supplementary Fig. S4B). This could reflect greater cellular density in young tissue, and increased levels of PTMs such as crosslinking in ageing, lessening detectability by MS. All matrisome classifications were represented in the analysis of F1 (Fig. 5B), with matrix regulators and matrix-affiliated proteins making up a greater proportion of the matrisome in young versus adult mice. In old and aged versus adult mice, proteoglycans made up a greater proportion of the matrisome at the expense of glycoproteins and matrix regulators.

**Figure 5.**
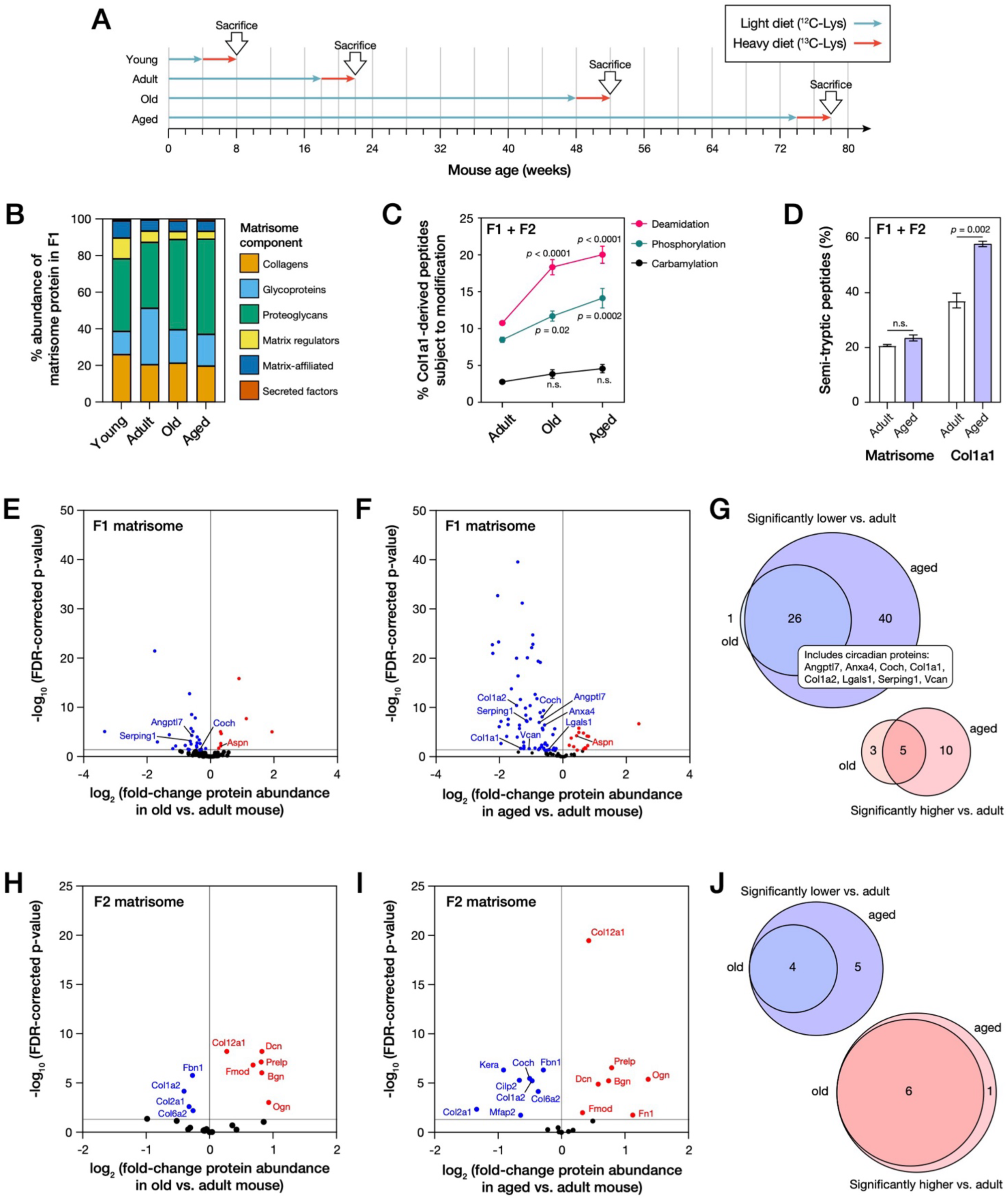
The protein composition of the mouse tail tendon is changed in ageing. (**A**) Schematic showing feeding regimes used in this study: mice were fed a ‘heavy diet’ (^13^C-lysine) for four weeks prior to sacrifice at age 8 weeks (‘young’ mouse group), 22 weeks (‘adult’), 52 weeks (‘old’), or 78 weeks (‘aged’). Following sacrifice, tail tendon tissue was fractionated into circadian/soluble (F1) and non-circadian/permanent (F2) fractions, as in earlier experiments. (**B**) Distribution of the total signal detected by mass spectrometry from matrisome proteins between the structure/function classifications of the matrisome in F1 of tail tendon tissue from young, adult, old and aged mice. Matrisome classifications are used as described by Shao *et al.* (46). (**C**) Average counts of Col1a1-derived peptides subjected to post-translational modification, expressed as percentages of the total number of Col1a1-derived peptides detected, in adult, old and aged mice; *p*-values of differences to adult from two-way ANOVA. (**D**) Average counts of matrisome- and Col1a1-derived semi-tryptic peptides, expressed as percentages of the total number of peptides derived from the matrisome and Col1a1, respectively, in adult and aged mice; *p*-values from t-tests. In parts (C) and (D), bars represent means ± s.e.m.; ‘n.s.’ = not significant. (**E**) Volcano plot showing fold-changes in matrix protein abundance in the F1 fraction of tail tendon from old vs. adult mice. (**F**) Volcano plot showing fold-changes in matrix protein abundance in the F1 fraction of tail tendon from aged vs. adult mice. (**G**) Venn diagrams illustrating the overlap between the matrix proteins down-regulated (blue) and up-regulated (red) in F1 fractions of tail tendon from old and aged mice. Matrix proteins found at lower abundance in both old and aged mice, relative to adult, included many that were previously identified as having significant circadian rhythmicity (Fig. 1H): Angptl7, Anxa4, Coch, Col1a1, Col1a2, Lgals1, Serping1 and Vcan (11). (**H**) Volcano plot showing fold-changes in matrix protein abundance in F2 from old vs. adult mice. (**I**) Volcano plot showing fold-changes in matrix protein abundance in F2 from aged vs. adult mice. (**J**) Venn diagrams illustrating the overlap between the matrix proteins down-regulated (blue) and up-regulated (red) in F2 fractions of tail tendon from old and aged mice. In figure panels (E), (F), (H) and (I), blue and red points indicate significant down- and up-regulation, respectively, with Benjamini-Hochberg false discovery rate (FDR) corrected *p*-values < 0.05. In (E) and (F), significantly changing matrisome proteins with circadian rhythmicity have been annotated (11); in (H) and (I), all significantly changing matrisome proteins are annotated. All data derived from proteomics analysis with *n* = 3 animals per age group.

To investigate the extent of PTMs in ageing in greater detail, we again focused on Col1a1 and widened our MS search parameters to capture deamidation, carbamylation, phosphorylation and semi-tryptic peptides. We found modifications to Col1a1 to be increased in old and aged mice, relative to adult (Fig. 5C). Consistent with our findings, deamidation and carbamylation are thought to occur increasingly with age, contributing to the accumulation of damage in tissue (54, 55). Increased phosphorylation of Col1a1 with ageing was unexpected. However, peptides detected in the old and aged mice were phosphorylated at a position (Ser535) previously associated with disruption to the triple helical structure of collagen (56). When collagen is denatured, this site is a substrate for extracellular signal-related protein kinase-1 (Erk1). This suggests that the type-I collagen helix may be disrupted in ageing and therefore more accessible to modification or cleavage. The aged mouse showed significantly more semi-tryptic Col1a1-derived peptides than the adult mouse, indicative of increased levels of degradation or fragmentation (*p* < 0.02; Fig. 5D).

### The circadian/soluble extracellular matrix pool is diminished in the tendons of ageing mice

We next considered age-associated changes in the abundances of specific matrisome components. Reassuringly, differences in the F1 matrisome in old versus adult mice were broadly reproduced and amplified in a comparison of aged versus adult (Figs. 5E, F). The majority of significant changes related to downregulation of proteins in the F1 pool: a common set of 26 proteins were significantly lower in both old and aged mice, versus adult (*p* < 0.05; Fig. 5G). This set included circadian-regulated matrisome components angiopoietin-related protein 7 (Angptl7), annexin A4 (Anxa4), cochlin (Coch), Col1a1, Col1a2, galectin-1 (Lgals1), plasma protease C1 inhibitor (Serping1), and versican (Vcan). This result demonstrates a suppression of circadian-regulated ECM in ageing, with potential consequences on tissue homeostasis and maintenance (57). Interestingly, the circadian proteoglycan Aspn counters this trend, exhibiting greater abundance in ageing. We also found commonality in old and aged versus adult changes in the F2 matrisome (Figs. 5H-J). Together, these findings are consistent with a characterisation of ECM ageing occurring as continuous, progressive decline, rather than as a rapid onset of failure (58). Proteins significantly increased in F2 in the old and aged mice included a number of small leucine rich proteoglycans (SLRPS): biglycan (Bgn), decorin (Dcn), prolargin (Prelp), fibromodulin (Fmod), and mimecan (Ogn); Fbn1, associated with mechanical load-bearing capacity, decreased in ageing.

A comparison of young versus adult F1 matrisome didn’t show a concerted loss or gain of circadian-regulated components (Supplementary Fig. S4C). The young F2 matrisome had lower levels of Coch, Prelp, Col12a1, Bgn, and CAP-Gly domain-containing linker protein 2 (Clip2), suggesting an ECM that was not yet fully matured (Supplementary Fig. S4D). Protein-lysine 6-oxidase (lysyl oxidase, lox), an extracellular enzyme that forms precursors to ECM crosslinks, was higher in young animals, perhaps required for ECM remodelling during maturation. Overall, continuous, monotonic changes between young and aged animals were not apparent, suggesting distinct processes of development and ageing.

### Protein turnover slows in ageing and the distinction between circadian/soluble and non-circadian/permanent pools of extracellular matrix is diminished

An examination of H/L ratios in all proteins in both F1 and F2 fractions revealed a significant decrease in heavy-label incorporation in the tendons of older mice (*p* < 0.0001; Supplementary Fig. S5A), consistent with slowing metabolism (26). GO analysis showed intracellular proteins with slowed turnover to be associated with biological processes of ATP metabolism, cell adhesion, and response to mechanical stimulation (Supplementary Fig. S5B). Increased half-lives of ECM components were determined across matrisome classes (Supplementary Figs. S5C-E), suggestive of increased crosslinking or decreased enzymatic degradation. Half-lives of matrisome proteins were substantially shorter in young mice, but in all age groups collagens remained significantly longer lived than glycoproteins (*p* < 0.01). As exemplified earlier by type-I collagens, we found a significant difference between the half-lives of matrisome proteins in F1 versus F2 pools of adult tendon tissue (*p* < 0.0001; Fig. 6A). However, a separation between matrisome half-lives in F1 versus F2 was not found in young mice (Fig. 6B). Our conjecture is that this was because young mice had not yet deposited their mature matrix, so both pools were subject to remodelling. Indeed, only three matrisome proteins in the F2 fraction of young mouse tendon (Col7a1, Col25a1, and Itih5) met our criteria for inclusion in a ‘permanent matrix’ (i.e., having half-lives exceeding one year; Fig. 3C). This was further reflected in universally lower half-lives of individual matrisome proteins in young versus adult mice in both F1 and F2 (Supplementary Figs. S5F, G).

**Figure 6.**
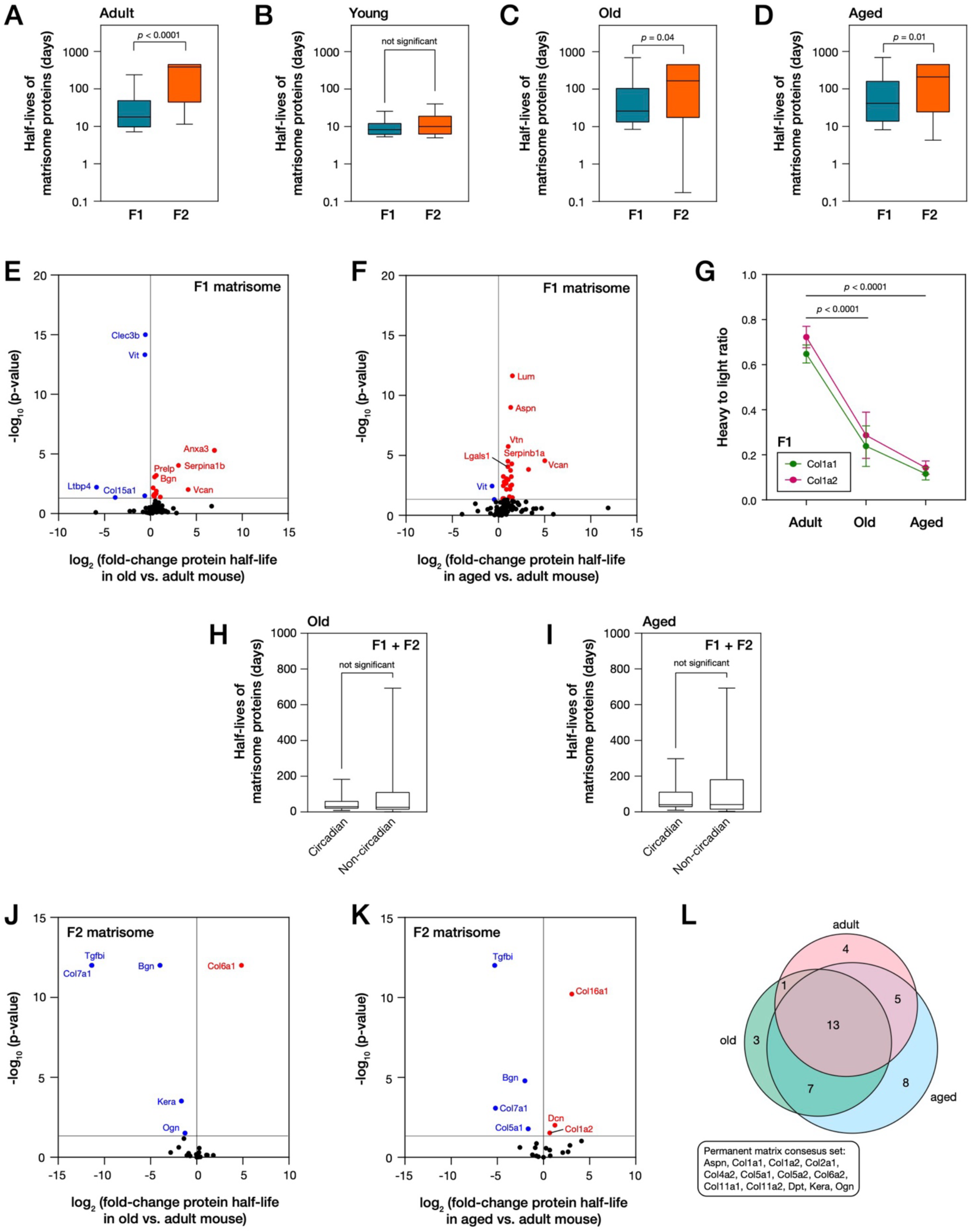
Older mice have slower matrix turnover in tail tendon, and distinctive properties of matrix pools are diminished. (**A**) Matrisome protein half-lives in F1 and F2 fractions of tail tendon from adult 22 week) mice. (**B**) Matrisome protein half-lives in F1 and F2 fractions of tail tendon from young (8 week) mice. (**C**) Matrisome protein half-lives in F1 and F2 fractions of tail tendon from old (52 week) mice. (**D**) Matrisome protein half-lives in F1 and F2 fractions of tail tendon from aged (78 week) mice. (**E**) Volcano plot showing fold-changes in matrisome protein half-lives in F1 fraction of tail tendon from old vs. adult mice. (**F**) Volcano plot showing fold-changes in matrisome protein half-lives in F1 fraction of tail tendon from aged vs. adult mice. (**G**) Average ratios of signal from heavy and light peptides from Col1a1 and Col1a2 in analysis of F1 fraction of tail tendon from adult, old and aged mice. The observance of significantly shorter half-lives of circadian vs. non-circadian proteins observed in adult mouse tendon (Fig. 2H) was not replicated in (**H**) old or (**I**) aged mice. (**J**) Volcano plot showing fold-changes in matrisome protein half-lives in F2 fraction of tail tendon from old vs. adult mice. (**K**) Volcano plot showing fold-changes in matrisome protein half-lives in F2 fraction of tail tendon from aged vs. adult mice. (**L**) Venn diagram comparing the permanent matrices (components with half-lives longer than one year) in adult, old and aged mice. In panels (A)-(D), (H) and (I), box-whisker plots indicate medians, 10^th^ and 90^th^ percentiles, minimum and maximum values; *p*-values determined by Mann-Whitney tests. In panels (E), (F), (J) and (K), blue and red points indicate proteins with significantly shortened and lengthened half-lives, respectively, with *p*-values < 0.05. In panel (G), points indicate mean ± s.e.m.; *p*-values from two-way ANOVA. All data derived from proteomics analysis with *n* = 3 animals per age group; protein half-lives determined using models described by Alevra *et al.* (49).

Contrastingly, in the older mice, whilst a significant difference between the half-lives of F1 and F2 remained (*p* < 0.04), the magnitude of the difference was decreased relative to the adult mouse (Figs. 6C, D). This narrowing separation largely resulted from increased half-lives of proteins in the F1 fraction (Figs. 6E, F), suggesting a loss of regulation on the faster turnover circadian matrix pool. This included the regulation of type-I collagens in F1, with both Col1a1 and Col1a2 incorporating significantly lower levels of the heavy label in old and aged mice versus adult (*p* < 0.0001; Fig. 6G). These observations are consistent with lost capacity for translation and protein metabolism in ageing (59), and suppression of circadian rhythmicity (57). The collagen-specific molecular chaperone 47 kDa heat shock protein (Serpinh1) showed faster turnover in F1 in old and aged mice. Serpinh1 expression has been found to decrease in ageing (60), perhaps indicative of misbalance between its synthesis and degradation, with downstream consequences on collagen regulation.

Hierarchical clustering of H/L ratios allowed us to identify age-associated trends in the F1 matrisome (Supplementary Fig. S6A). The analysis identified four main behaviours in turnover rates in adult, old and aged mice: late-onset decrease; steady decrease; early-onset increase; and late-onset increase (Supplementary Figs. S6B-E). The majority of the matrisome was placed into clusters where protein turnover slowed with age; importantly, these clusters also contained all quantified components of the circadian-regulated tendon matrisome (Angptl7, Anxa4, Col1a1, Col1a2, Coch, Lgals1, Lgals3, and Vcan) (11). Furthermore, where adult mice showed a significant difference between half-lives of circadian and non-circadian matrisome components (*p* = 0.03, Fig. 2H), this distinction was absent in old and aged mice (Figs. 6H, I). Together, these results indicate a suppression of circadian matrix regulation in ageing tendon. A corresponding analysis of changes to individual matrisome protein half-lives and hierarchical clustering in F2 again showed a general slowing of protein turnover (Figs. 6J, K; Supplementary Fig. S6F). Constituents of the permanent matrix established to persist for longer than one year in the F2 pool (Fig. 3C) were generally maintained in old and aged mice (Fig. 6L). However, the size of the permanent matrix set increased with age, including incorporation of basement membrane laminin proteins, suggestive of an age-associated failure of turnover.

## DISCUSSION

### The presence of co-existing matrix pools with distinct rates of turnover

In this study we have separately analysed circadian/soluble (F1) and non-circadian/permanent (F2) pools of ECM from mouse tail tendon. Type-I collagen present in both these pools had significantly different half-lives: 4 ± 2 days in F1 compared to 700 ± 100 days in F2. Our analysis supports and provides mechanistic insight to the model proposed by Chang *et al.* in which a ‘lifelong’ pool of collagen is complemented by a second, shorter-lived pool, which is maintained over a circadian cycle (11). This pattern of ECM regulation goes beyond type-I collagen, and we have identified multiple rapidly remodelled proteins, as well as an interacting set of ‘permanent matrix’ proteins. Our approach measured the half-lives of 1325 unique proteins, with an average half-life of 63 days. Previous analysis has shown the average half-lives of proteins in cartilage tissues to vary from 8 -11 days, and those in muscle 10-14 days (43), but it is plausible that fractionation of these tissues would reveal greater diversity in rates of ECM turnover. Our results suggest that matrix homeostasis is maintained by a complex, multi-component mechanism. A requirement for both F1 and F2 matrix pools in tendon can be rationalised by the former being sacrificial and acting to protect the permanent structure. Circadian rhythmicity of the matrix has been indicated in tissues such as cartilage (61), intervertebral disc (20), as well as in other tendons. We might therefore propose that the presence of co-existing but separately regulated matrix pools could be a systemic mechanism for homeostasis in ECM-rich tissues subjected to circadian cycles of stress and strain.

### Dysregulation of matrix turnover in ageing

Loss of matrix homeostasis, tissue weakness and increased susceptibility to injury are commonly observed in ageing tissues concomitantly with a dampening of circadian rhythmicity (57). Our data supports the notion that age-associated loss of circadian rhythmicity leading to disruption of the balance between fast and slow turnover ECM pools. This disruption could lead to loss of homeostasis, and therefore to compromised integrity and capacity for remodelling and repair. Daily rhythmic patterns in mechanical loading of the tendon (i.e., corresponding to differences in activity levels between day and night), in addition to mechanical abnormalities observed in the tendons of circadian-knockout mice, suggest that the fast-turnover circadian/soluble ECM pool plays a mechanical role in the tissue (11). A reduced ability of aged mice to respond to mechanical loading has been associated with decreased collagen biosynthesis as well as reduced matrix metalloproteinase (MMP) activity (62). This would suggest a corresponding change to ECM turnover, consistent with the findings of our study. Dysregulated turnover of a protective ECM pool could contribute to the slowing of the healing process observed in aged mice (63). Moreover, as premature aging is observed in mice with collagenase-resistant type-I collagen, turnover of the circadian/soluble pool appears important for matrix longevity (32).

Type-I collagens are the most prevalent in tendon tissue and showed the most striking differences in turnover rate between F1 and F2 pools, but our analysis was able to capture additional regulatory patterns within the matrisome. Many of these were also affected by ageing. The set of interacting permanent matrix proteins – those with half-lives of a year or longer – that were identified in adult mice were not yet established in immature mice. The permanent matrix was maintained in aged mice, but contained additional glycoproteins such as laminins and the tenascin (Tnc). This suggests a decline in protein remodelling machinery, or an increase in crosslinking causing aberrant persistence in ECM proteins that would be turned over in younger animals. The age-associated decline in the mechanical properties of the tendon have been proposed to be linked to increased type-V collagen, which is a regulator of collagen fibril assembly (27). Our data support this, showing faster turnover of collagen-V with age. Ageing caused a decrease in the turnover rate of Bgn in F2, but an increase in F1. Dcn, another proteoglycan important in tendon matrix assembly, showed contrasting faster turnover in F2 with age. Mice lacking Bgn or Dcn have been shown to have abnormal collagen fibrils with modified mechanical properties (34). Together, this evidence suggests that balanced proteoglycan turnover is required for ECM homeostasis, but that this is dysregulated in ageing.

Aside from synthesis and degradation, PTMs can affect turnover and are prevalent in the ECM. Phosphorylation has been found to influence protein turnover using a similar pulse-labelling approach to that deployed here (64). PTMs such as AGEs and crosslinking have been shown to accumulate in tendon with age, decreasing protease accessibility and therefore degradation. This would likely have a larger effect on the faster turnover F1 pool. Moreover, circadian regulation of the protease cathepsin K (Ctsk) has been predicted to play a role in rhythmic collagen levels observed in tendon (11). Changes in F1 fibril accessibility to Ctsk could therefore explain the observed reduction in turnover in aged tendon. A similar explanation could also answer the question of how these smaller fibrils are selectively turned over, rather than the larger fibrils found in the permanent matrix. The matrix is also heavily glycosylated, including glycosaminoglycan (GAG) side chains which coat collagen fibrils. These side chains are required for Ctsk cleavage and so this could also impact turnover (65). Understanding the specific mechanisms which separate the fast and slow turnover pools, and how these are affected in ageing, could allow therapeutic intervention to maintain homeostasis and integrity in ECM-rich tissues.

### Matrix turnover as a potential target for biomarker discovery

Dampened circadian rhythms in tendinopathy patients, combined with reduced type-I collagen synthesis at night, implicate the tendon circadian clock as a therapeutic target (15). Applied to the data presented here, this suggests altered dynamics of the faster turnover F1 pool in tendon inflammation/damage. Monitoring this pool, as well as its turnover, could therefore provide valuable biomarkers of tendinopathy. Furthermore, the altered dynamics of this pool in ageing suggest a clear link to the pathology observed in tendon inflammation. Investigations into protein turnover have previously been connected to disease signatures and biomarkers and are thus emerging as a valuable research area within clinical settings (35, 44). This is further indicated by increased use of biomarkers of collagen synthesis or degradation in enzyme-linked immunosorbent assay (ELISA)-based screens of patient serum samples where these indicators of turnover have been implicated in diseases such as arthritis (66), fibrosis (67) and cancer (68).

## MATERIALS AND METHODS

### Animal work

The care and use of all animals in this study were carried out in accordance with the UK Home Office regulations, UK Animals (Scientific Procedures) Act of 1986 under Home Office Licences (#70/8858 or I045CA465). For circadian time-course studies, wild-type C57BL/6 mice housed in a cycle of 12 hour light and 12 hour dark were harvested every 4 hours across 48 hours. For heavy-isotope labelling studies, male wild-type C57BL/6 mice were fed on a heavy-lysine diet (^13^C-Lys; CK Isotopes) for 1, 2, 4 or 8 weeks, prior to culling at an age of 8, 22, 52 or 78 weeks. Control mice were fed an equivalent normal-lysine diet (CK Isotopes). Tail tendon tissue was dissected for analysis as described by Chang *et al.* (11).

### Preparation of mouse tail tendon tissues for label-free analysis

Analysis of the proteins extracted from mouse tail tendon tissue over a circadian time course using a mild detergent wash (fraction 1, ‘F1’) was reported by Chang *et al.* (11). In this study, a subsequent high salt extraction protocol (fraction 2, ‘F2’) was applied to the same material. Pellets were suspended in 75 µL High Salt buffer (3 M sodium chloride, NaCl, Fisher Scientific; 25 mM dithiothreitol, DTT, Sigma; 25 mM ammonium bicarbonate, AB, Sigma Aldrich; protease and phosphatase inhibitor cocktail in accordance with the manufacturer’s guidance, Sigma) with 6 x 1.6 mm steel beads (Next Advance) and homogenised for 5 minutes using a bullet blender (Next Advance) at maximum speed. Samples were then centrifuged (20 k g for 5 minutes) and supernatants (F2) were transferred to LoBind Protein tubes (Eppendorf). Protein concentrations were determined using bicinchoninic acid (BCA) assay (BCA Gold, Thermo, according to the manufacturer’s protocol).

### Preparation of mouse tail tendon tissue for analysis following heavy-isotope labelling

200 µL SL-DOC buffer (1.1% sodium dodecanoate, Sigma; 0.3% sodium deoxycholate, Sigma; 25 mM AB; 0.5 mM DTT; protease and phosphatase inhibitor cocktail in accordance with the manufacturer’s guidance, Sigma) was added to the mouse tail tendon tissue, in addition to 6 x 1.6 mm steel beads (Next Advance). Samples were then homogenised for 5 minutes using a bullet blender (Next Advance) at maximum speed. 12 µL iodoacetamide (IAA, Sigma) was then added to 30 mM, and the samples left in darkness for 20 minutes before quenching with 12 µL 30 mM DTT. Samples were then centrifuged (20 k g for 5 minutes) and the supernatants (F1) were transferred to LoBind Protein tubes. The pellets were then further solubilised in 75 µL High Salt buffer, subjected to an additional 5 minutes in the bullet blender at maximum speed, centrifuged (20 k g for 5 minutes), and the supernatants transferred to fresh LoBind Protein tubes. The remaining pellets were then transferred to 500 µL Covaris tubes, and resuspended in 350 µL Lysis buffer (5% sodium dodecyl sulphate, SDS, Sigma; 50 mM tetraethylammonium bromide, TEAB, Sigma). Samples were then disrupted in a Covaris LE220+ sonicator using a 40 W lysis programme with 100 cycles per burst, a duty factor of 40% and a peak incident power of 500 W. Samples were then reduced with 19 µL DTT (final concentration, 5 mM) at 60 °C, alkylated with 33 µL IAA (final concentration, 45 mM), and quenched with 57 µL 0.1 M DTT. Following centrifugation (20 k g for 5 minutes), the supernatants from the High Salt and Lysis buffer extractions were pooled to give F2. Protein concentrations in F1 and F2 were determined by BCA assay (BCA Gold, Thermo, according to the manufacturer’s protocol).

### Preparation of samples for mass spectrometry analysis

50 µg of protein from each sample was prepared in 5% SDS to make a total volume of 50 µL, and then acidified using 5 µL of 12% H_3_PO_4_. 350 µL binding buffer (90% methanol, 10% distilled water; 100 mM TEAB) was added and the samples loaded onto S-Trap columns (ProtiFi) according to the manufacturer’s protocol. Column-bound proteins were washed in Binding buffer and methyl tert-butyl ether (MTBE, Sigma), and then digested with 5 µg of trypsin (Promega, from 0.8 µg/µL stock solution) in 65 µL Digestion buffer (50 mM TEAB; 0.1% formic acid, FA, Sigma; at pH 8.5). Peptides were eluted from the column in 65 µL Digestion buffer, 65 µL 0.1% FA in distilled water, and finally with 30 µL 0.1% FA, 30% acetonitrile (ACN, Sigma) in water. Peptides were desalted using Oligo R3 resin beads (Thermo Scientific), according to the manufacturer’s protocol, in a 96-well, 0.2 µm polyvinylidene fluoride (PVDF) filter plate (Corning). The immobilised peptides were washed twice with 0.1% FA prior to elution in 2x 50 µL 0.1% FA with 30% ACN, and lyophilisation using a speed-vac (Heto Cooling System).

### Mass spectrometry

Peptides were resuspended in 10 µL 0.1% FA in 5% ACN. Liquid chromatography (LC) separation was performed on a Thermo RSLC system consisting of an NCP3200RS nano pump, WPS3000TPS autosampler and TCC3000RS column oven configured with buffer A as 0.1% FA in water and buffer B as 0.1% DA in ACN. The analytical column (Waters nanoEase M/Z Peptide CSH C18 Column, 130 Å, 1.7 µm, 75 µm x 250 mm) was kept at 35 °C and at a flow rate of 300 nL/minute for 8 minutes. The injection valve was set to load before a separation consisting of a 105 minute multistage gradient ranging from 2 to 65% of buffer B. The LC system was coupled to a Thermo Exploris 480 mass spectrometry system via a Thermo Nanospray Flex ion source. The nanospray voltage was set at 1900 V and the ion transfer tube temperature set to 275 °C. Data was acquired in a data-dependent manner using a fixed cycle time of 2 seconds, an expected peak width of 15 seconds and a default charge state of +2. Full mass spectra were acquired in positive mode over a scan range of 300 to 1750 Th, with a resolution of 120000, a normalised automatic gain control (AGC) target of 300% and a max fill time of 25 ms for a single microscan. Fragmentation data was obtained from signals with a charge state of +2 or +3 with an intensity over 5000 and they were dynamically excluded from further analysis for a period of 15 seconds after a single acquisition within a 10 ppm window. Fragmentation spectra were acquired with a resolution of 15000, a normalised collision energy of 30%, an AGC target of 300%, a first mass of 110 Th, and a maximum fill time of 25 ms for a single microscan. All data was collected in profile mode.

### Proteomics data analysis

Mass spectrometry data was analysed using Proteome Discoverer (Thermo Fisher). Datasets from heavy-isotope labelling experiments were analysed using a SILAC 1plex (Lys6) method. All searches included the fixed modification for carbamidomethylation on cysteine residues resulting from IAA treatment. The variable modifications included in all analyses were oxidation of methionine (monoisotopic mass change, +15.955 Da) and phosphorylation of threonine, serine and tyrosine (+79.966 Da). A maximum of 2 missed cleavages per peptide was allowed. The minimum precursor mass was set to 350 Da with a maximum of 5000 Da. Precursor mass tolerance was set to 10 ppm, fragment mass tolerance was 0.02 Da and minimum peptide length was 6. During specific analysis of post translational modifications, additional variable modifications associated with the ECM were included in the MS search. These included: oxidation/hydroxylation of aspartic acid, phenylalanine, histidine, lysine, asparagine, arginine, tryptophan and tyrosine (+15.955 Da); phosphorylation of aspartic acid and histidine; deamidation of asparagine and glutamine (+0.984 Da); carbamylation of lysine (+43.006 Da). These searches were also semi-tryptic to allow detection of degraded proteins.

Peptides were searched against the SwissProt database using the Sequest-HT search engine with a maximum false discovery rate of 1%. Proteins were required to have a minimum false discovery rate (FDR) of 1% and at least 2 unique peptides in order to be accepted, known contaminants were also removed. Missing values were assumed to be due to low abundance. Abundance comparisons in aged mice were done using MaxQuant (v1.6.14) using default parameters with the exception of multiplicity being set to 2 and the heavy lysine label being added; peptide modifications were set as described above. Abundances were compared using MsqRob where LFQ data was normalised using median peptide intensity (69, 70).

Heavy-to-light ratios were determined using the abundance of a protein detected in the ‘heavy’ channel, containing ^13^C-lysine, compared to the ‘light’ channel. Heavy-to-light ratios were converted to half-lives using code developed by Alevra *et al.* (49), with a minimum of 5 data points required for half-life calculation. Further statistical analysis and data visualisation was performed using R, including MsqRob, MetaCycle and Pheatmap packages (version 4.1.2), and Prism (GraphPad Software, version 9.1.2). Statistics in Prism were analysed by two-way ANOVA or Mann-Whitney tests, unless otherwise stated; data is presented as mean ± s.e.m. Network analysis was performed using STRING (version 11.5) to generate protein interactions, filtered for high confidence scores, which were then exported into Cytoscape (version 3.8.2) for visualization. Gene ontology (GO) analysis was performed using DAVID2021, with proteins lists of interest being compared to a total protein background. Pathway enrichment was determined using FDR-corrected *p*-values.

### Periodicity analysis

Spectra acquired from mass spectrometry were aligned using Progenesis QI (Nonlinear dynamics) using default settings, except for peak picking which was set to 4/5. Only peptides with a charge between +1 and +4, and two-or-more isotopes were used in analysis. Mascot (Matrix Science) was used to search against SwissProt and TREMBL databases. The same modifications and settings were used as described previously and peptide and protein identifications were filtered as above. A mixed-effects linear regression model was fitted to proteins to calculate differences in protein quantity across the time points as previously described by Chang *et al.* (11). Periodicity was analysed in R using MetaCycle (71), default parameters were used and a Benjamini-Hochberg FDR <0.2 threshold was set to determine rhythmicity.

## ACKNOWLEDGEMENTS

The work was funded by the Biotechnology and Biological Sciences Research Council (BBSRC; BB/T001984/1), the Wellcome Trust (110126/Z/15/Z and 203128/Z/16/Z), the Medical Research Council (MRC; MR/P010709/1), and Arthritis Research UK (20875). AH was supported by a BBSRC CASE DTP studentship. JC was supported by an MRC Career Development Award (MR/W016796/1). JS was supported by a BBSRC David Phillips Fellowship (BB/L024551/1). Proteomics was performed by the Biological Mass Spectrometry Core Facility (supported by BBSRC, Wellcome and the University of Manchester Strategic Fund). We thank the University of Manchester Biological Support Facility for expertise and assistance in animal welfare and husbandry, and Drs Craig Lawless and Julian Selley for bioinformatics support.

## AUTHOR CONTRIBUTIONS

Conceptualization, AH, JC, QJM, KEK, and JS; Investigation, AH, JC, MFAC, RO’C, and SW; Formal Analysis, AH, JC, SW, QJM, KEK, and JS; Writing – Original Draft, AH; Writing – Visualization, Review & Editing, AH, JC, MFAC, RO’C, SW, DK, QJM, KEK, and JS; Project Administration and Funding Acquisition, JC, DK, QJM, KEK, and JS.

## DATA AVAILABILITY STATEMENT

Proteomics data have been deposited to the ProteomeXchange Consortium via the PRIDE partner repository with the identifier PXD042915.

