## Supplementary Figures for "Circadian and permanent pools of extracellular matrix co-exist in tendon tissue, but have distinct rates of turnover and differential responses to ageing"

(1) Wellcome Centre for Cell-Matrix Research, Oxford Road, Manchester, M13 9PT, UK.
(2) Division of Cell Matrix Biology and Regenerative Medicine, School of Biological Sciences, Faculty of Biology, Medicine and Health, Manchester Academic Health Science Centre, University of Manchester, Manchester, M13 9PL, UK.
(3) Current address: Kennedy Institute of Rheumatology, University of Oxford, Oxford, OX3 7FY, UK.
(4) Division of Molecular and Cellular Function, School of Biological Sciences, Faculty of Biology, Medicine and Health, Manchester Academic Health Science Centre, University of Manchester, Manchester, M13 9PL, UK.
(5) Current address: School of Chemistry, Chemical Engineering and Biotechnology, Nanyang Technological University, Singapore, 637459 Singapore.
(6) Biological Mass Spectrometry Core Facility, Faculty of Biology, Medicine and Health, Manchester Academic Health Science Centre, University of Manchester, Manchester, M13 9PL, UK.
(7) Current address: Inoviv, LABS Hogarth House, 136 High Holborn, London, WC1V 6PX, UK.

**CONTENTS**

Supplementary Figure S1 … … … … … … … … … 3

Supplementary Figure S2 … … … … … … … … … 4

Supplementary Figure S3 … … … … … … … … … 5

Supplementary Figure S4 … … … … … … … … … 6

Supplementary Figure S5 … … … … … … … … … 8

Supplementary Figure S6 … … … … … … … … … 10

References … … … … … … … … … 12

### SUPPLEMENTARY FIGURES

#### Supplementary Figure S1

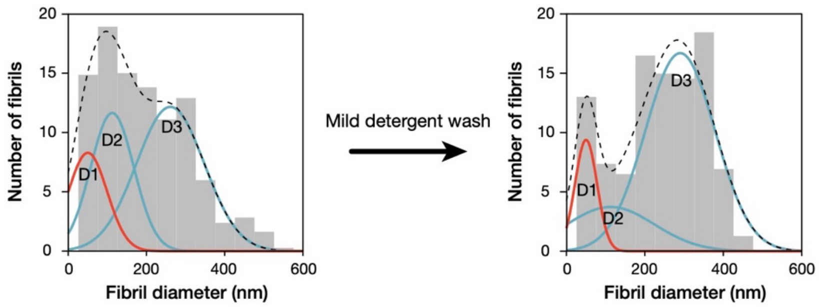


**Figure S1. Mild detergent wash of mouse tail tendon preferentially solubilises narrower fibrils.** Analysis of transmission electron microscopy (TEM) images published by Chang *et al.* (1) showing mouse tendon fibrils before and after a mild detergent wash, quantifying preferential removal of D1 fibrils (diameter < 75 nm) relative to D2 and D3 fibrils (diameters 75 – 150 nm and > 150 nm, respectively). Curve fits to the sum of three gaussian distributions performed in Igor Pro (Wavemetrics); methodology described by Starborg *et al.* (2).

**Supplementary Figure S2**

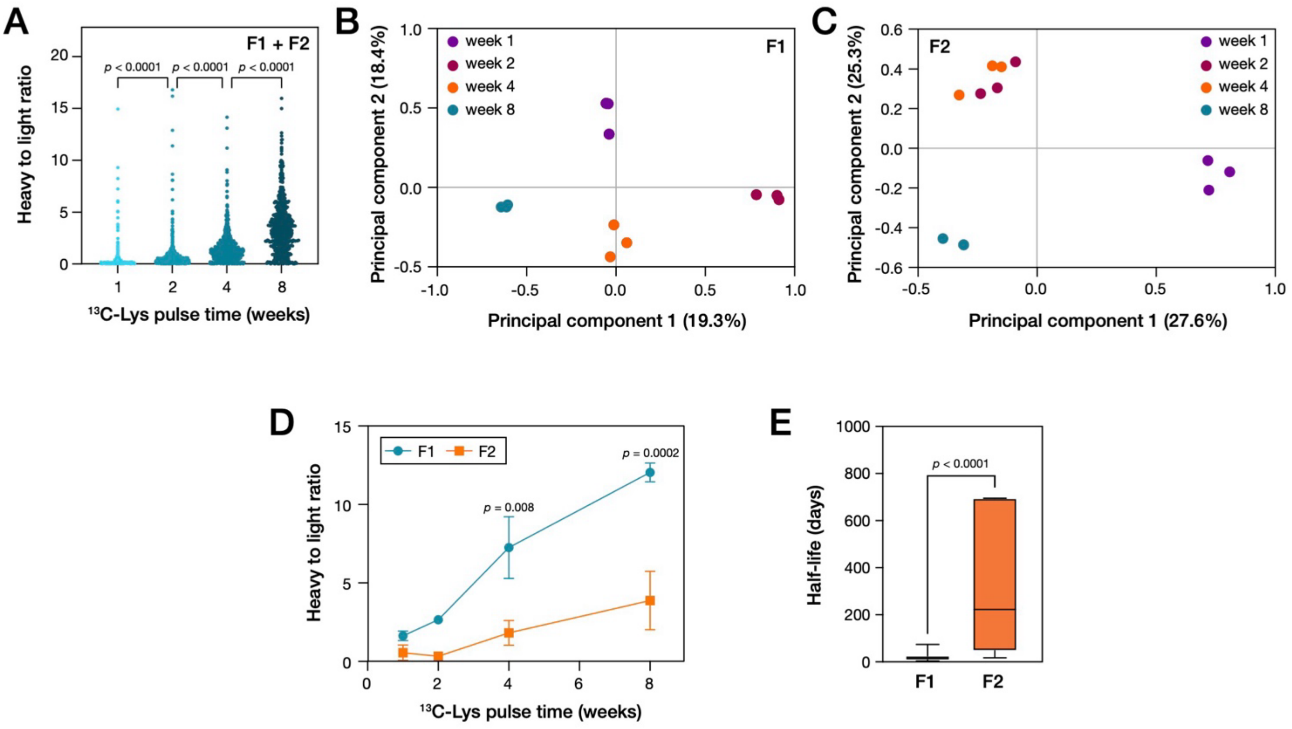


**Figure S2. Longer periods of feeding on the $^{13}$C-Lys diet caused greater heavy isotope incorporation in both F1 and F2 fractions.** (**A**) The distribution of ratios of heavy to light peptides derived from proteins in tail tendon increasingly favoured heavy as the mice were fed the $^{13}$C-Lys diet for longer periods. Plot combines data from both F1 and F2 fractions; *p*-values from Kruskal-Wallis tests. (**B**) Principal component analysis (PCA) plot of peptide-level proteomics data (F1 extraction from tail tendon at 22 weeks of age) following feeding on a $^{13}$C-Lys diet for 1, 2, 4 or 8 weeks. (**C**) Corresponding PCA plot for the F2 fraction. In both F1 and F2 data was found to cluster according to the length of time animals were fed the heavy isotope diet. (**D**) Ratios of signal from heavy and light peptides derived from all proteins in analysis of F1 and F2 fractions of tail tendon from 22 week old mice fed a $^{13}$C-Lys diet for 1, 2, 4 or 8 weeks. Points indicate mean ± s.e.m.; *p*-values from two-way ANOVA. (**E**) Protein half-lives were significantly longer in F2 vs. F1; *p*-value from Mann-Whitney test. Proteomics analysis from $n$ = 3 animals per condition; protein half-lives determined using models described by Alevra *et al.* (3).

**Supplementary Figure S3**

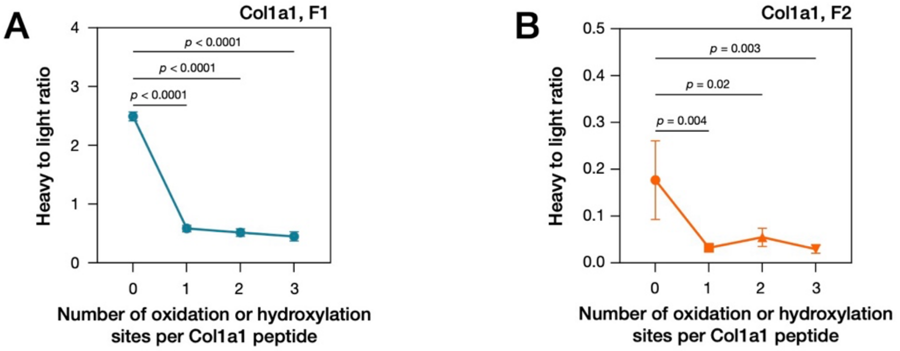


**Figure S3. Oxidation of Col1a1 peptides.** (**A**) Average ratios of signal from heavy and light peptides derived from Col1a1 in F1, in 22 week old mice fed a $^{13}$C-Lys diet for 4 weeks, as a function of the number of oxidation or hydroxylation modifications detected per peptide. (**B**) Average ratios of signal from heavy and light peptides derived from Col1a1 in F1, in 22 week old mice fed a $^{13}$C-Lys diet for 4 weeks, as a function of the number of oxidation or hydroxylation modifications detected per peptide. In both (A) and (B), significantly higher isotope labelling was found in unoxidized peptides; *p*-values from Kruskal-Wallis tests.

### Supplementary Figure S4

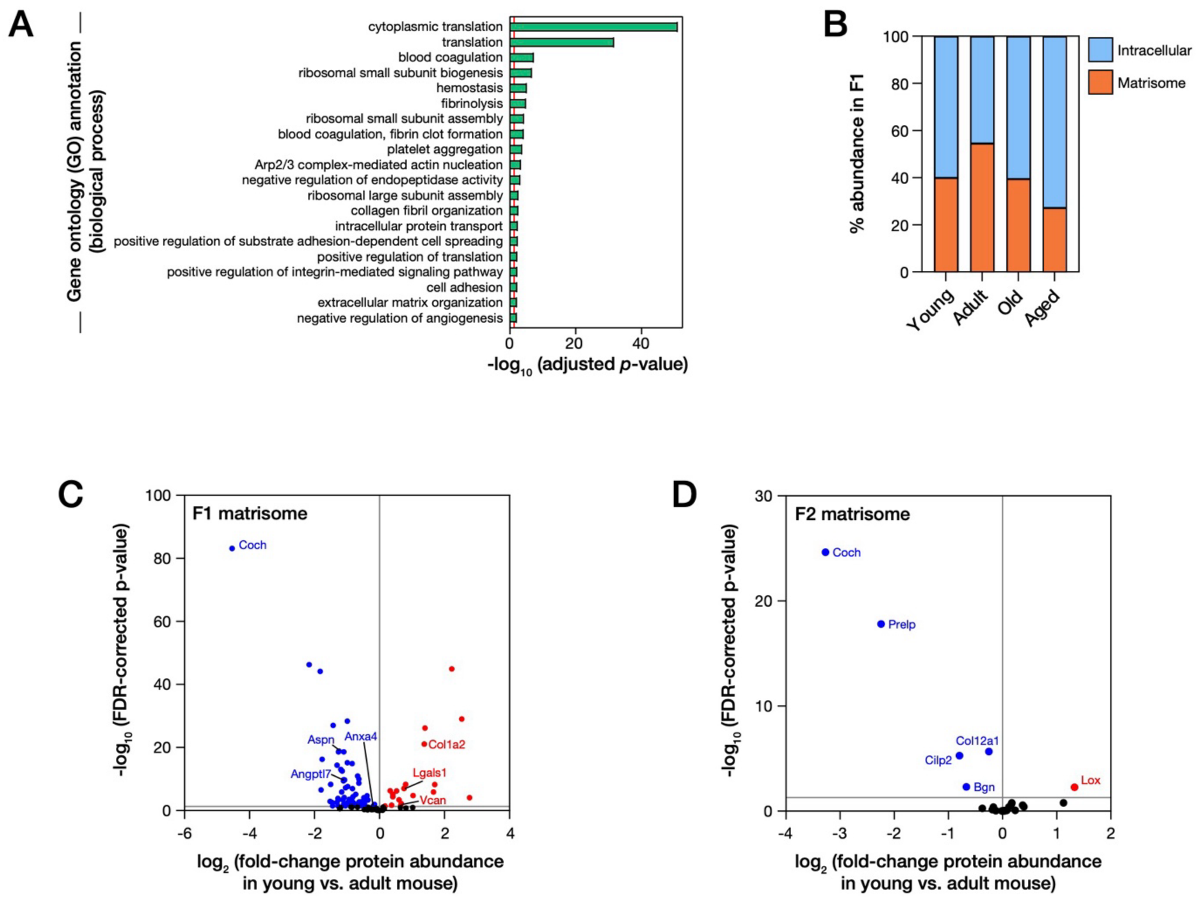

**Figure S4. Effect of ageing on the proteomic composition of mouse tail tendon.** Mice were fed a 13C-lysine diet for 4 week prior to sacrifice at 8 weeks ('young'), 22 weeks ('adult'), 52 weeks ('old') and 78 weeks ('aged'). Tail tendon tissue was excised, solubilized in F1 and F2 fractions, and analysed by mass spectrometry (MS). (**A**) Biological process gene ontology (GO) terms significantly enriched in the set of proteins downregulated ($\log_2$-fold change < -0.5) in both old and aged tendon tissue, relative to adult. (**B**) Fraction of the total signal detected by mass spectrometry from peptides derived of intracellular vs. matrisome proteins in F1 of tail tendon tissue from young, adult, old and aged mice. Matrisome components are described by Shao *et al.* (4). **C**) Volcano plot showing fold-changes in matrix protein abundance in the F1 fraction of tail tendon from young vs. adult mice. (**D**) Volcano plot showing fold-changes in matrix protein abundance in the F2 fraction of tail tendon from young vs. adult mice. In panels (C) and (D), blue and red points indicate significant down- and up-regulation, respectively, with Benjamini-Hochberg false discovery rate (FDR) corrected $p$-values $< 0.05$. In (C), significantly changing matrisome proteins with circadian rhythmicity have been annotated; in (D), all significantly changing matrisome proteins are annotated. All data derived from proteomics analysis with $n = 3$ animals per condition.

**Supplementary Figure S5**

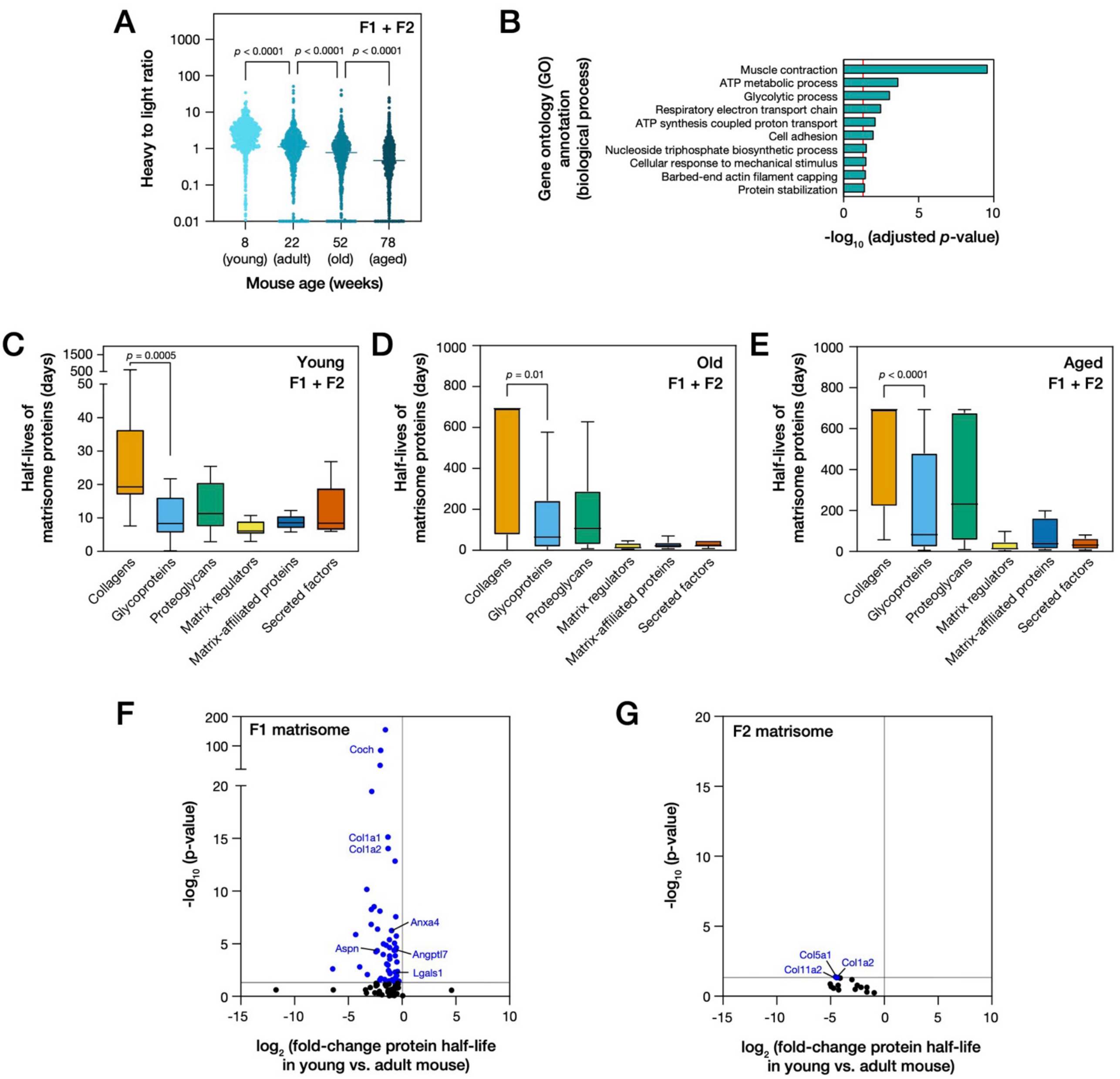

**Figure S5. Effect of ageing on protein turnover rates in mouse tail tendon.** (**A**) Distributions of ratios of heavy to light peptides derived from all proteins in tail tendon of young (sacrificed at 8 weeks), adult (22 weeks), old (52 weeks) and aged (78 weeks) mice. The rate of incorporation of the heavy isotope was significantly slowed with animal age. (**B**) Gene ontology (GO) analysis of biological process annotations enriched in the set of intracellular proteins with slower turnover ($\log_2$-fold change < -0.5) in both old and aged tendon tissue, relative to adult. (**C**) Distribution of half-lives of ECM proteins in the tail tendon of young mice within the matrisome structure/function classifications described by Shao *et al.* (34). (**D**) Distribution of half-lives of ECM proteins in the tail tendon of old mice. (**E**) Distribution of half-lives of ECM proteins in the tail tendon of aged mice. (**F**) Volcano plot showing fold-changes in matrisome protein half-lives in F1 fraction of tail tendon from young versus adult mice. (**G**) Volcano plot showing fold-changes in matrisome protein half-lives in F2 fraction of tail tendon from young versus adult mice. Analysis in (A), (C), (D) and (E) combines F1 and F2 data; box-whisker plots indicate medians, $10^{th}$ and $90^{th}$ percentiles, minimum and maximum values; *p*-values from Kruskal-Wallis tests. In panels (F) and (G), blue points indicate proteins with significantly shorter half-lives, with $p$-values < 0.05. All data derived from proteomics analysis with $n = 3$ animals per condition; protein half-lives determined using models described by Alevra *et al.* (3).

### Supplementary Figure S6

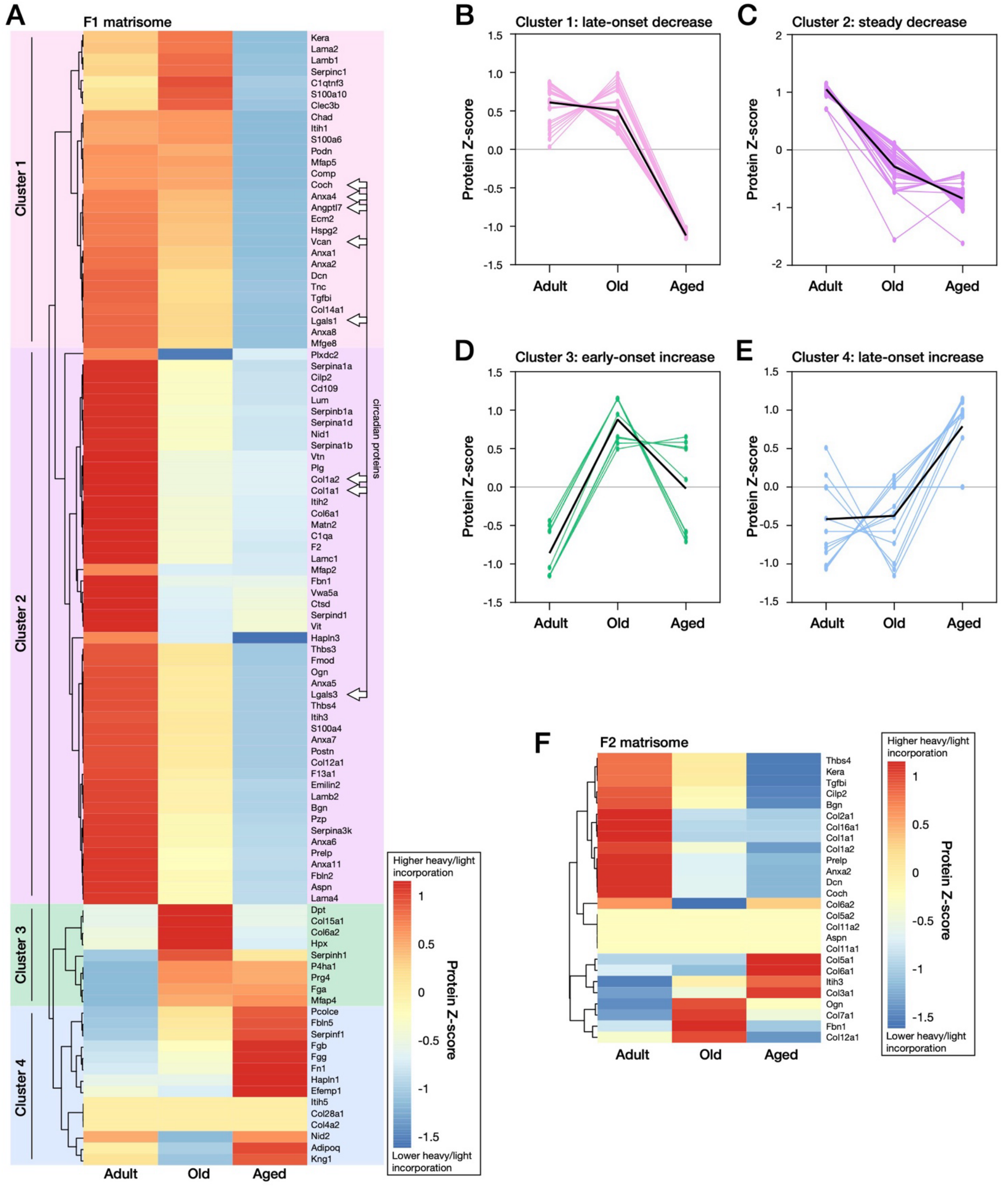

**Figure S6. Hierarchical clustering analysis of changes in matrisome turnover in tail tendons of ageing mice.** (**A**) Hierarchical clustering of turnover rates of matrisome proteins in F1 of tail tendon from adult (sacrificed at 22 weeks), old (52 weeks) and aged (78 weeks) mice. Proteins were assigned into one of four groups by k-means clustering. Proteins identified as having circadian rhythmicity in the soluble F1 fraction (1) were found in clusters 1 and 2. (**B**) Proteins in cluster 1 exhibited a late-onset decrease in turnover rate (i.e., in aged mice only). (**C**) Cluster 2 (inclusive of Col1a1 and Col1a2) exhibited a steady decrease in turnover rate. (**D**) Cluster 3 exhibited an early-onset increase in turnover rate (i.e., manifesting both in the old and aged mice). (**E**) Cluster 4 exhibited a late-onset increase in turnover rate. (**F**) Hierarchical clustering of turnover rates of matrisome proteins in F2 of tail tendon from adult, old and aged mice. Data derived from proteomics analysis of mouse tail tendon with $n$ = 3 animals per condition; protein half-lives determined using models described by Alevra *et al.* (3).
